# A two-scale model of Legionnaires’ disease to predict incubation periods and risk of symptomatic disease

**DOI:** 10.64898/2026.01.22.701076

**Authors:** Nyall Jamieson, Christiana Charalambous, David M. Schultz, Ian Hall

## Abstract

*Legionella pneumophila* is an intracellular pathogen that causes Legionnaires’ disease, a severe pneumonia acquired primarily through contaminated water systems. Public health interventions rely on accurate estimates of the incubation period and dose–response (DR) relationship, yet currently used approaches in the literature underestimate incubation periods by assuming Markovian rupture times for infected macrophages. Here, we develop the first non-Markovian, two-scale within-host framework for Legionnaires’ disease, coupling stochastic intracellular replication in individual macrophages with extracellular Legionella–macrophage population dynamics. At the cellular level, we model intracellular replication using a stochastic logistic birth–death (SLBD) process, coupled with non-Markovian rupture-time distributions (Erlang and Burr). The Erlang distribution preserves tractability via the method of separation, whereas the Burr distribution captures heavy-tailed rupture times consistent with experimental data. Simulations are implemented using a renewal-based non-Markovian Gillespie algorithm. At the host level, successive infection and rupture events describe population-scale infection dynamics, enabling estimation of DR curves and incubation-period distributions. Across six model variants, DR predictions remain robust, with ID_50_ estimates narrowly ranging between 8.79 and 8.94 *Legionella*, consistent with guinea pig challenge data. In contrast, incubation-period estimates show strong dependence on rupture-time assumptions: non-Markovian models predict median incubation periods of 5–6 days, correcting the previous 2–3 day underestimation and aligning with human outbreak data (2–10 days, up to 13 days). Sensitivity analysis identifies rupture size, phagocytosis rates, and threshold effects as key determinants of incubation-period results. By relaxing exponential assumptions, our framework provides biologically realistic within-host dynamics that improve epidemiological predictions. These results refine the quantitative basis for outbreak investigations and environmental risk assessment and are generalizable to other intracellular pathogens such as *Coxiella burnetii* and *Francisella tularensis*.

**Author summary:** *Legionella pneumophila* causes Legionnaires’ disease, a serious pneumonia often linked to contaminated water systems, but key quantities such as the incubation period remain difficult to estimate accurately. Existing models assume that infected immune cells rupture at random times with no memory, an assumption that simplifies mathematics but does not reflect experimental observations. We developed a model that follows bacterial growth inside individual macrophages and connects these cellular events to infection dynamics within a host. Unlike previous approaches, our model allows rupture times to follow more realistic, non-exponential patterns that better match laboratory data. Using simulations, we show that commonly used assumptions systematically underestimate the incubation period of Legionnaires’ disease. Our results predict incubation periods of 5–6 days, consistent with human outbreak data, while leaving estimates of infectious dose largely unchanged. This work improves the biological realism of within-host infection models and provides a stronger quantitative foundation for outbreak investigation and environmental risk assessment. The modelling framework can be adapted to study other intracellular pathogens that replicate inside host immune cells.

## 1 Introduction

*Legionella pneumophila* is a gram-negative bacterium that causes legionellosis [1]. This disease manifests as either severe pneumonia (Legionnaires’ disease) or a milder respiratory illness (Pontiac fever) [2]. Legionnaires’ disease develops when aerosolised *Legionella* are inhaled into the lungs. Upon inhalation, *Legionella* infect alveolar macrophages and replicate intracellularly. After a latent period, infected host cells rupture, releasing the *Legionella* progeny population to infect additional macrophages. This cycle either ends by all *Legionella* dying or disease progression, which results in symptoms within an individual. These symptoms range from fever and cough to respiratory failure and, in severe cases, death [1, 3]. Unlike many respiratory diseases, Legionnaires’ disease rarely transmits person-to-person; infection risk arises primarily from contaminated water systems [3].

Because transmission occurs via environmental exposure, public health investigations depend on models linking infection biology to epidemiological outcomes. Two key quantities underpin these investigations: the incubation-period distribution [4] and the dose–response (DR) relationship [5], which defines the probability of illness given an exposure dose. Inaccurate incubation-period estimates hinder public health officials from correctly identifying exposure windows, reducing the likelihood of finding the source of contamination and leaving populations at increased risk [6]. Likewise, a mischaracterized DR relationship prevents the establishment of effective exposure limits or cleaning protocols in water systems, potentially resulting in preventable infections.

Mathematical models of *Legionella* and other intracellular pathogens, including *Francisella tularensis* and *Coxiella burnetii*, frequently employ continuous-time Markov chains (CTMCs) with exponentially distributed rupture times [7–10] in an attempt to estimate the incubation period and DR of these diseases. Although exponential waiting times simplify model implementation, they poorly reflect biological reality: rupture probability of macrophages infected with *Legionella* exhibit a sigmoidal increase as a function of time [7, 11]. Markovian approximations reduce model flexibility and accuracy, leading to underestimated incubation periods, thereby amplifying the public health consequences described above.

To address this limitation, we develop a non-Markovian, two-scale within-host framework for Legionnaires’ disease. At the cellular scale, each infected macrophage follows a stochastic logistic birth–death (SLBD) process until rupture, generating a time-dependent rupture-size distribution once combined with non-Markovian rupture-time distributions. Specifically, we model rupture times using two distributional families: the Erlang and Burr [12] distributions. The Erlang distribution, embedded in CTMCs via the method of stages, preserves analytical tractability while capturing median rupture times [13–15]. On the other hand, the Burr distribution, which fits experimental data most accurately, captures both heavy-tailed behavior and median dynamics [12]. Therefore, the Erlang and Burr distributions offer a compromise between analytical convenience and biological realism. Because Burr distributions cannot be represented in finite-state CTMCs, we implement the non-Markovian Gillespie algorithm (nMGA), a renewal-based extension that tracks event ages and accommodates general hazard functions [16]. At the host scale, extracellular bacteria infect new macrophages, which undergo independent intracellular dynamics, producing a stochastic process of infection, replication, and macrophage rupture until clearance or symptom onset. These approaches allow us to evaluate both tractable approximations and biologically flexible models.

We quantify model outputs by simulating both the Erlang CTMC and Burr nMGA models, with parameterizations informed by human and guinea pig data [11, 17–19]. We then validate the framework against known experimental results [20]. Our approach may also be applied to other intracellular pathogens, including *Coxiella burnetii* and *Francisella tularensis*, offering comparison to prior CTMC and phase-type models [9, 10, 21–23]. Specifically, some models have assumed purely a CTMC approach [9, 10], with no consideration for non-Markovian rupture times or time dependency on the rupture size. Extensions have been developed for Tularensis and Anthrax that preserve a CTMC approach but with a more complex phase-type distribution [21, 23].

To our knowledge, this paper provides the first non-Markovian, two-scale within-host model of Legionnaires’ disease. By combining Erlang compartmentalisation and Burr-based nMGA simulations with an explicit stochastic model of intracellular replication, our framework extends CTMC approaches to incorporate biologically realistic rupture-time distributions. Specifically, relaxing exponential assumptions corrects the previous underestimation, producing incubation period and low-dose risk predictions consistent with experimental and outbreak data. This extension demonstrates the critical role of reliably representing intracellular dynamics in accurately modelling epidemiological quantities such as the incubation period of Legionnaires’ disease.

## 2 Materials and methods

In this section, we describe the biological dynamics that occur post-infection with *Legionella*, as well as the details of the previous within-host model of Legionnaires’ disease described in [7]. In addition to this, we outline the assumptions used here to extend this model and derive a novel within-host model of Legionnaires’ disease. Section 2(a) begins by describing the internal processes that occur once *Legionella* infects an individual. Following this, Section 2(b) presents the model from our previous work [7], which we will refer to as the *CTMC model*. Next, Section 2(c) presents the parameter estimates used in the CTMC model and our new model. Section 2(d) discusses the deterministic within-macrophage model and exponential rupture-time model defined in the CTMC model. We include here a derivation of our stochastic extension to the within-macrophage model as well as our non-Markovian rupture-time distribution extensions. Then, to conclude our Methods section, Section 2(e) discusses our two-scale within-host models, which incorporate the rupture-size distribution that is obtained from modelling with our stochastic within-macrophage model and new rupture-time models.

### 2.1 Within-host dynamics of *Legionella* infection

We consider the biological scenario presented in [7], which describes infection following a single inhalation of aerosols containing *Legionella*. Upon exposure, individuals may inhale varying numbers of bacteria, some of which survive and establish infection in the lungs. Alveolar macrophages, as the main phagocytic cells within the lungs, attempt to eliminate bacteria through phagocytosis. During phagocytosis, a macrophage either destroys the bacterium or fails, in which case intracellular replication of *Legionella* occurs. The bacterial population grows inside the macrophage until the macrophage ruptures, releasing *Legionella* back into the lung environment. Cytokines released during these processes recruit additional immune cells to the lungs, including neutrophils, monocytes, and dendritic cells. These other phagocytes are typically recruited at least five days after human infection [24, 25]. By this time, infection has usually progressed to either recovery or symptom onset. For our within-host model, we focus only on alveolar macrophages, as they are the principal phagocytes in *Legionella* infection [24]. This choice simplifies parameter estimation to one cell type.

Estimates place the total number of alveolar macrophages in both lungs at about 10^9^ [26]. Given this large population, we approximate the lungs as containing infinitely many macrophages. This assumption allows a simplified linear model of infection dynamics. Additionally, this assumption treats the macrophage population as roughly constant during infection, which reduces analytical complexity. We assume that each macrophage engulfs at most one *Legionella* at a time. Thus, a macrophage either eliminates the bacterium or becomes infected and eventually ruptures. Further, because extracellular bacteria are sparse relative to macrophages, we assume that at most one bacterium infects a macrophage. Furthermore, in vivo, inflammation from cytokine responses to phagocytosis and macrophage rupture drives symptom onset. Therefore, for simplicity, we approximate inflammation levels with the extracellular *Legionella* population and assume that symptoms appear once the extracellular *Legionella* population exceeds a threshold *T*_*L*_.

### 2.2 Continuous-time Markov chain (CTMC) model

We introduce the CTMC model developed in [7] to describe the biological process once *Legionella* deposit in the lungs. This model used a two-dimensional Markov chain (*L*(*t*), *M* (*t*)), with *L*(*t*) and *M* (*t*) representing the number of extracellular *Legionella* and infected macrophages at time *t*. Three key events occur during *Legionella* infection: a macrophage engulfs and kills an extracellular *Legionella*, an extracellular *Legionella* infects a macrophage, or an infected macrophage ruptures and releases the new intracellular *Legionella* population. The CTMC contains two absorbing states at *t*_absorb_ > 0: recovery [(*L*(*t*_absorb_), *M* (*t*_absorb_)) = (0, 0)] or symptom onset [*L*(*t*_absorb_) = *T*_*L*_].

*α* was defined as the rate at which an individual extracellular *Legionella* survives phagocytosis, *β* was defined as the rate at which an individual extracellular *Legionella* is killed from phagocytosis, and *λ* was defined as the rupture rate of an individual infected macrophage. Thus, *α* + *β* gave the total phagocytosis rate of individual extracellular *Legionella*. These events were modelled as Markovian random variables for parsimony. Furthermore, two infected-macrophage rupture scenarios were considered. In the first scenario, each rupture releases *G* bacteria as in [9, 10]. In the second scenario, rupture sizes followed a negative binomial distribution with mean *G* and dispersion parameter *r*, extending [9, 10]. The state-change rates in the latter case were:

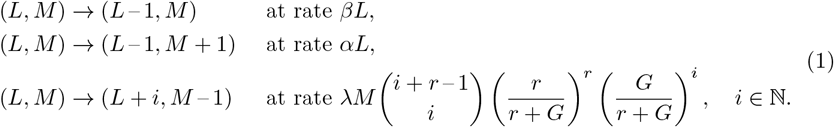

Alternatively, the final rate is replaced with *λM* when assuming each rupture releases exactly *G* bacteria [9, 10]. Three within-host models were developed using this CTMC framework. *Model A* assumed a fixed rupture size of *G Legionella*, which occurred at rate *λL. Model B* assumed a negative binomial rupture-size distribution with mean *G* and dispersion parameter *r*. This rupture event also occurred at rate *λL*. Finally, *model C* assumed a bootstrapped rupture-size distribution, which was obtained by sampling parameter estimates for the CTMC model and propagating the uncertainty. Time-independent rupture-size distributions were assumed across all three models. Additionally, the Gillespie algorithm [27] was used to simulate the three within-host models until either absorbing state occurred and repeated simulations for doses from 1 to 500. 1000 simulations per dose were conducted to estimate probability of illness.

### 2.3 Model parameter estimates

For context, we briefly summarize the parameter estimates used in the CTMC model [12], as well as the new model we develop in this paper. These parameters define the infected-macrophage rupture rate used in the CTMC model (*λ*), rupture size (*G*), rate of successful phagocytosis (*α*), rate of failed phagocytosis (*β*), the threshold population for symptom onset (*T*_*L*_), intracellular bacterial growth rate (*ω*), and intracellular carrying capacity (*ψ*). Parameter values were derived from experimental datasets of macrophage rupture times, intracellular bacterial counts, and extracellular bacterial growth in mice and guinea pigs [11, 17, 18]. Distributions for each parameter were obtained via bootstrapping to reflect uncertainty in the estimates.

**Table 1.**
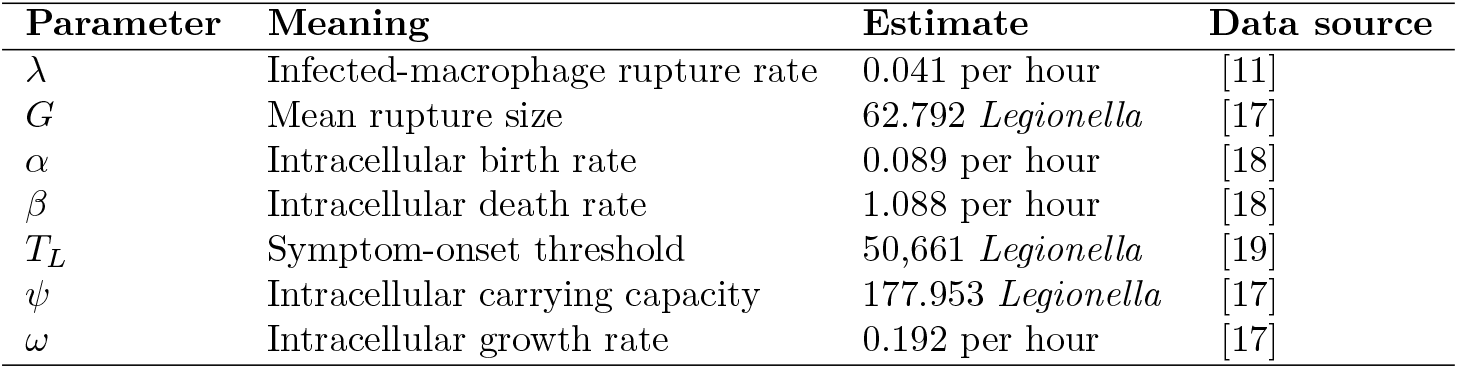
Summary of parameter estimates derived in the previous within-host CTMC model [7].

| Parameter | Meaning | Estimate | Data source |
| --- | --- | --- | --- |
| $\lambda$ | Infected-macrophage rupture rate | 0.041 per hour | [11] |
| $G$ | Mean rupture size | 62.792 *Legionella* | [17] |
| $\alpha$ | Intracellular birth rate | 0.089 per hour | [18] |
| $\beta$ | Intracellular death rate | 1.088 per hour | [18] |
| $T_L$ | Symptom-onset threshold | 50,661 *Legionella* | [19] |
| $\psi$ | Intracellular carrying capacity | 177.953 *Legionella* | [17] |
| $\omega$ | Intracellular growth rate | 0.192 per hour | [17] |

### 2.4 Within-macrophage models

A hybrid model of infected macrophage dynamics was developed as part of the CTMC model. This hybrid model combined an exponential distribution for the rupture-time process with a deterministic intracellular bacterial logistic-growth model. These processes were separated to allow independent modelling. In this section, we first describe the rupture-time distribution of the CTMC model and then develop our extension for the rupture-time model. Following this, we describe the deterministic within-macrophage model and then develop our stochastic extension of this model.

#### 2.4.1 Rupture-time distribution

To model rupture timing, the exponential, Erlang, and Burr distributions have previously been fitted to experimental data for rupture times of bone-marrow–derived macrophages infected with *Legionella* [11]. In this work, the Burr distribution provided the best fit to the experimental data, whilst the exponential distribution fitted poorly (Figure 1). Specifically, the exponential distribution underestimated the mean and overestimated the variance when fitted to data. However, the exponential distribution was chosen to preserve the Markov property, which allowed for simple implementation as a CTMC model.

**Fig 1.**
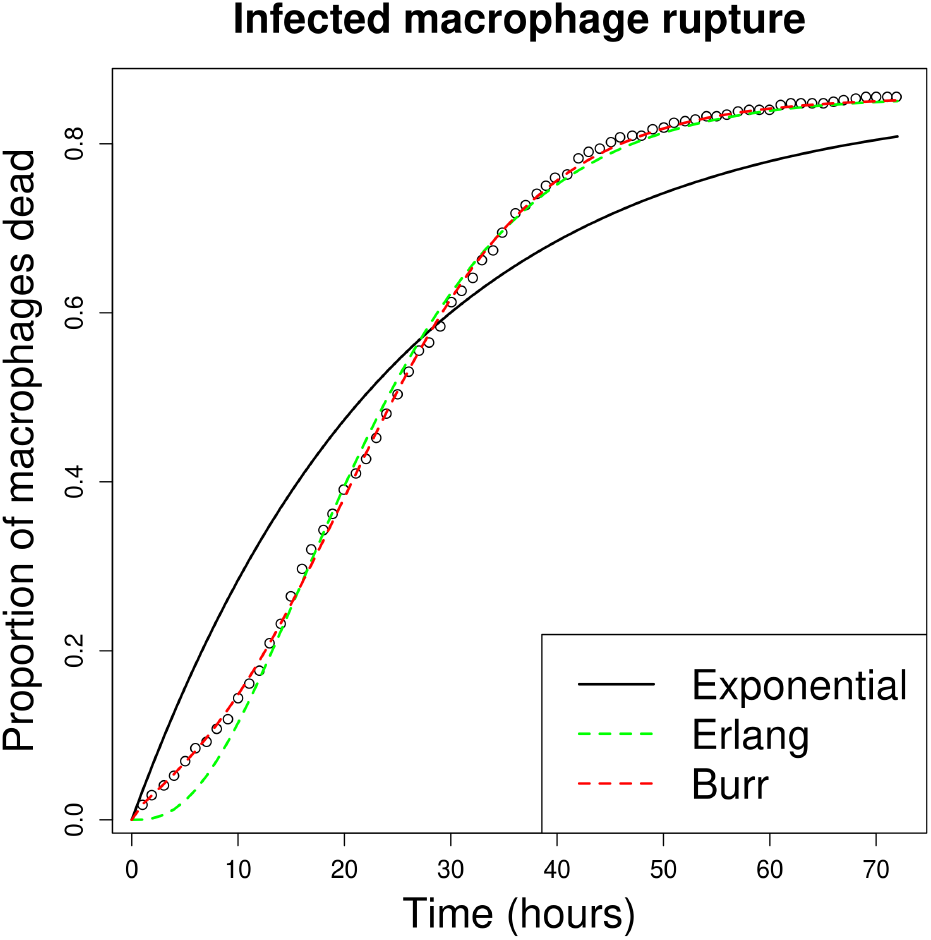
Time series of the proportion of bone-marrow-derived macrophages that were killed by Legionella hourly between 1–72 hours [11]. The three time-delay distributions were fitted to the data.

For our rupture-time model, we employ both the Erlang and Burr distributions in two distinct approaches, as these distributions provide a better fit to the data [11]. We define *T* as a random variable that represents the rupture time of an infected macrophage. Under the Erlang model *T* ∼ Erlang(*α*_*E*_, *β*_*E*_), the probability density function is:

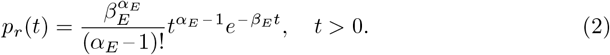

Fitting this distribution to the rupture-timing data, we obtained 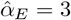 and 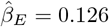 (s.e. 7.486 × 10 ^−4^). For the Burr model we have the following probability density function [12]:

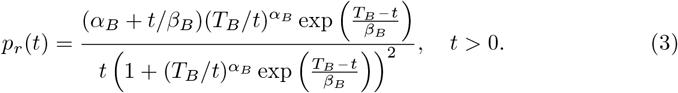

Fitting to the experimental data, this model gave 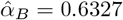 (s.e. 0.04671), 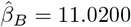 (s.e. 0.27457), and 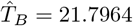 (s.e. 0.06471).

#### 2.4.2 Within-macrophage birth–death model

For the CTMC model, a deterministic logistic-growth process was defined for the intracellular-growth process. This model captured initial exponential replication of intracellular *Legionella*, followed by a reduced growth rate due to intracellular resource constraints and crowding. These assumptions aligned with observed macrophage infection dynamics [17], and were defined as follows:

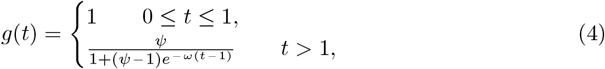

where *ψ* represents the intracellular carrying capacity of *Legionella* and *ω* represents the intracellular growth rate of *Legionella*. This equation represented the mean growth of the *Legionella* population within a macrophage at time *t* after infection. A rupture-size distribution was derived with three different approaches. First, for model A, *G* was fixed at the median rupture time *g*(1*/λ*). Second, for model B, *λ, ψ* and *ω* were sampled from their bootstrapped distributions to compute a sample of *G* by calculating *g*(1*/λ*). A negative binomial rupture-size distribution to this sample. In models A and B, all other parameters remained fixed at their point estimate. Third, for model C, the above bootstrapped sample of *G* was used. In model C, they allowed variation for all parameters by sampling new estimates between iterations of model simulation. In all three scenarios, they assumed independence between the rupture-time and rupture-size models. This independence assumption preserved the Markovian framework and avoided explicitly modelling the time-since-infection component.

Here, we extend the above intracellular *Legionella* growth process. Specifically, we construct a SLBD process that is a stochastic analogue of equation (4). To model stochastic intracellular dynamics, we use a master equation approach. This approach calculates the full time-dependent distribution of intracellular bacterial counts instead of relying on simulated approximations. We define the probability distribution *p*_*l*_(*t*) as the probability that the number of *Legionella* remaining alive in a macrophage at time *t* after infection is *l*. We may define a vector *p*(*t*) = (*p*_1_(*t*), *p*_2_(*t*), …)^*T*^ and a transition rate matrix *Q* such that:

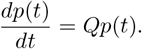

Here, the *i*^th^ row and *j*^th^ column of *Q* provide the rate at which the intracellular *Legionella* count transitions from *i* to *j* in a single birth–death event. We construct the master equations from single-step transition probabilities over a time interval *δt*. These probabilities capture bacterial birth and death rates. For a state *l* ∈ ℕ, if *l* > 1, birth occurs in (*t, t* + *δt*) with probability *ωlδt* + *o*(*δt*), whereas death occurs with probability *ωl*^2^*δt/ψ* + *o*(*δt*). If *l* = 1, birth occurs in (*t, t* + *δt*) with probability *ωl*(1 – *l/ψ*)*δt* + *o*(*δt*). Using these transition probabilities, we obtain:

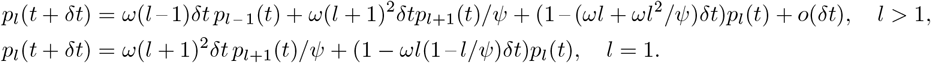

These differential equations yield the continuous-time master equation for intracellular bacterial counts:

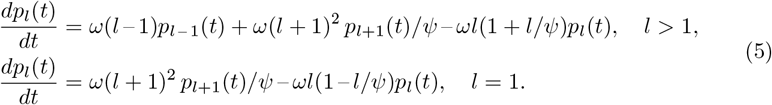

#### 2.4.3 Combining rupture-time and intracellular-*Legionella* growth models

Now that both the rupture-time and intracellular-*Legionella* growth models have been defined, we combine these two types of models to derive time-dependent rupture-size distributions. In the literature, combining these two components typically involved considering birth–death–killing (or catastrophe theory) [21–23]. In these examples, analytical solutions were considered for the rupture-size distributions and the rupture-time distribution when assumed that the hazard rate for a rupture event is proportional to the intracellular load [21, 22]. However, we consider a simpler approach and propose three approaches to link the delay distributions with the SLBD model. First, we consider the equation:

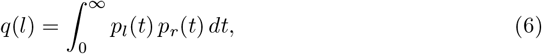

where *p*_*l*_(*t*) represents the intracellular population distribution at a time *t* since infection, and *p*_*r*_(*t*) is the empirical rupture-time distribution. Here, *q*(*l*) is defined to represent the probability that a number of *l Legionella* are released upon infected-macrophage rupture.

As visualised in Figure 1, the empirical cumulative distribution function of macrophage rupture times is *defective* (i.e., it does not converge to one for large *t*) [11]. Consequently, a subset of macrophages remains unruptured; however, the underlying cause cannot be identified from the data alone. Since our modelling approach for linking rupture timing and bacterial load is data-driven rather than mechanistic, we do not attempt to infer the reasons why some macrophages never rupture. Instead, we treat the rupture-time and intracellular growth datasets as independent characterizations of distinct aspects of the population dynamics. Directly sampling from equation (6) could therefore introduce bias by implicitly assuming a mechanism for non-rupturing macrophages that is not supported by the data.

To avoid this, we marginalize equation (6) to obtain a low-order approximation, the *average* rupture-size distribution

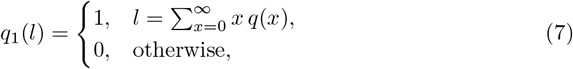

which captures the typical burst behaviour across macrophages while avoiding bias from unresolvable mechanisms. In this way, the population model is driven by reproducible, data-fit dynamics without imposing additional mechanistic structure where the data are inconclusive.

To reintroduce variability lost in this low-order approximation, we incorporate heterogeneity at the macrophage level by resampling the within-macrophage parameters *ω* and *ψ* across simulations. This approach provides a distribution rather than a point mass *q*_1_(*l*). Thus, the *intermediate* rupture-size distribution *q*_2_(*l*) is mathematically defined by equations (6) and (7), but with natural variability included via heterogeneous parameter sampling. This captures realistic macrophage-to-macrophage variation while remaining consistent with the data-driven, low-order approximation philosophy.

Third, we approximate variability around the point-estimate *q*_1_(*l*) using a Poisson rupture-size distribution. This *Poisson* distribution has mean equal to the point mass in (7), and is defined as follows:

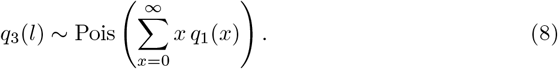

Although both rupture timing and intracellular replication are modelled stochastically at the within-cell scale, the rupture-size distribution used at the population scale only needs to summarise this variability through its expected value. The Poisson distribution provides a parsimonious approximation in which the mean of the mechanistic growth model directly specifies the stochastic rupture size, without introducing additional parameters beyond those already encoded in the intracellular model. In this sense, Poisson acts as a low-parameter summary of variability already generated by the mechanistic dynamics, rather than an additional source of biological uncertainty.

These models [*q*_1_(*l*), *q*_2_(*l*), and *q*_3_(*l*)] represent the three rupture-size models in our two-scale framework. They differ in complexity. The average model, *q*_1_(*l*), preserves a description of average rupture-size dynamics across a homogeneous macrophage population. The intermediate model, *q*_2_(*l*), preserves a description of average rupture-size dynamics, with macrophage-to-macrophage heterogeneity for variability. The Poisson model, *q*_3_(*l*), approximates variability using a simple Poisson distribution. All three models were derived from assuming dependence of the rupture timing on the intracellular *Legionella* population size.

### 2.5 Within-host model

Next, we extend the CTMC framework by incorporating alternative rupture-time distributions *p*_*r*_(*t*) and linking them to rupture-size models *q*_1_(*l*), *q*_2_(*l*) and *q*_3_(*l*). Specifically, we derive two families of within-host model: a model with the Erlang rupture-time distribution for *p*_*r*_(*t*) defined in equation (2), and another model with the Burr-distributed rupture-time distribution for *p*_*r*_(*t*) defined in equation (3). The Erlang-based model preserves the Markov property by separating compartments [13–15]. This approach enables simulation using the standard Gillespie algorithm. However, the Burr-based model introduces non-Markovian inter-event times. Therefore, we simulate this model using the nMGA to account for time-dependent hazard rates, with the within-host extensions derived below.

#### 2.5.1 Erlang-distributed infected-macrophage lifespan

We derive an extended CTMC model that results from replacing an exponential rupture-time distribution with an Erlang distribution. While Erlang-based compartmental extensions of CTMC frameworks have been used previously in population-level and epidemiological settings [13–15], to our knowledge this approach has not been applied to within-host bacterial growth and rupture dynamics. Existing within-host models typically assume exponential event timing, which can be overly simplistic, or adopt more complex phase-type formulations, which may be unnecessarily cumbersome; here, we show use an Erlang formulation as a parsimonious yet accurate alternative that balances simplicity and biological realism. The Erlang distribution is particularly well suited to this setting because it can be expressed as the convolution of *α*_*E*_ exponentially distributed random variables. Specifically, for our Erlang model, if *T* ∼ Erlang(*α*_*E*_, *β*_*E*_, then *T* ∼ Exp(*β*_*E*_) + … + Exp(*β*_*E*_) where the sum consists of *α*_*E*_ independent and identically distributed exponential random variables. Therefore, using the Erlang distribution in the compartmental model defined in equation (2) is equivalent to splitting the infected compartment into three sequential stages, capturing more realistic variability than a single exponential while avoiding the complexity of full phase-type models.

Specifically, we interpret the Erlang shape parameter *α*_*E*_ = 3 as dividing the infected-macrophage lifespan into *early, middle*, and *late* stages. Mathematically, we split the compartment *M* into three compartments (*M*_*E*_, *M*_*M*_, *M*_*L*_). Further, because the rupture-time data shows a defective probability distribution, we scale the final-stage Erlang rate by *θ* = 0.855, which is the limit at which the proportion of macrophages dying tends towards for large time, as suggested in Figure 1. This adjustment modifies the final rupture rate to *θβ*_*E*_ to reflect the proportion of macrophages that rupture. The staging method produces a compartmental model in which the infected-macrophage population progresses sequentially through early, middle, and late compartments. Each compartment is governed by Erlang-distributed rupture times, with the state transition rates defined as follows:

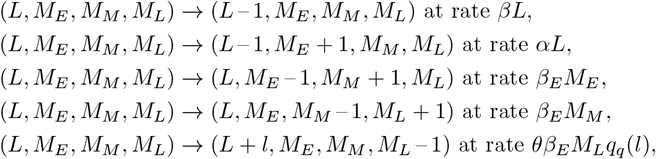

for *q* ∈ { 1, 2, 3 }. We simulate the model using the Gillespie algorithm to generate DR and incubation-period results.

#### 2.5.2 Burr-distributed infected-macrophage lifespan

We extend the within-host model in [7] by adopting a Burr distribution for rupture times. We simulate Burr-distributed rupture times using the nMGA [16]. This method accommodates non-exponential inter-event times in stochastic simulations. In this case, the Gillespie algorithm generalizes in this framework to handle non-Markovian inter-arrival times.

Again, we retain the assumptions that extracellular *Legionella* undergo phagocytosis and either survive at rate *αL* or die at rate *βL*, with both events following exponential distributions. For clarity, we define three event types: (i) surviving phagocytosis and infecting a macrophage (*i* = 1), (ii) dying from phagocytosis (*i* = 2), and (iii) infected-macrophage rupture (*i* = 3). Let *t*_*i*_ denote the elapsed time since the last occurrence of process *i*. The conditional probability density that the next event of type *i* occurs at time *τ* + *t*_*i*_, given that no such event has occurred before time *t*_*i*_ is:

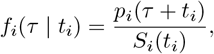

where *p*_*i*_ gives the probability density of process *i* and *S*_*i*_ its corresponding survival probability. Specifically, *p*_1,2_ are exponential probability density functions with rates *αL* and *βL* respectively. Additionally, *p*_3_ follows either the Erlang or Burr probability density function *p*_*r*_(*t*) defined in Section 2(d)(i). To simulate a statistically correct sequence of events, we define the joint density–mass function Θ(*τ, i* | {*t*_1_, *t*_2_, *t*_3_}), with *τ* > 0 continuous and *i* ∈ {1, 2, 3} discrete. Given the elapsed times {*t*_1_, *t*_2_, *t*_3_} for each event type, the joint density–mass function Θ(*τ, i* | { *t*_1_, *t*_2_, *t*_3_ }) specifies that the next event corresponds to process *i* and occurs at time *t* + *τ* . Therefore, since the conditional probability that a process *k* ≠ *i* does not occur in the the interval [*t, t* + *τ* ] is *S*_*k*_(*τ* | *t*_*k*_) = *S*_*k*_(*τ* + *t*_*k*_)*/S*_*k*_(*t*_*k*_), we obtain

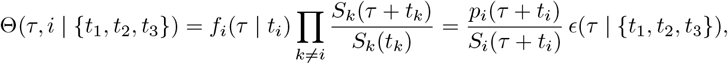

where

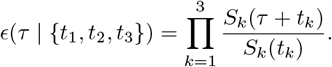

Given occurrence time *τ*, the probability that the next event belongs to process *i* is defined as

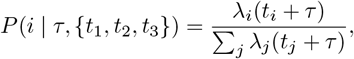

where the instantaneous hazard rate is defined as

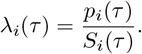

Therefore, the nMGA algorithm proceeds as follows:

1. Initialise elapsed times {*t*_*i*_}.
2. Draw a random waiting time *τ* by solving

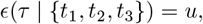

where *u* ∼ *U* (0, 1), and update global time *t* ← *t* + *τ* .
3. Select the next event type *i* from the distribution

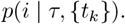
4. Update elapsed times: *t*_*k*_ ← *t*_*k*_ + *τ* for all *k* ≠ *i*, and reset *t*_*i*_ = 0.
5. Update the system state and active processes. If a new process becomes active, initialise its elapsed time.
6. Return to step (ii) and repeat until the simulation terminates.

### 2.6 Summary

We derived two rupture-time distributions and three rupture-size distributions. The rupture-size distributions (average, Poisson, and intermediate) differed in complexity, but all preserve the dependence between rupture timing and the intracellular bacterial population at rupture. Additionally, our two developed rupture-time distributions (Erlang and Burr) allow for more accuracy in the timing of rupture events. Combining all models yields six two-scale within-host models for Legionnaires’ disease. We simulate each model with an initial deposited dose of *L* = 1, …, 500, and 1000 iterations are conducted for each dose. Each simulation produces a binary outcome: symptom onset or recovery. However, in symptomatic cases, the simulation records the incubation-period estimate. We analyze the DR relationship and dose-dependent incubation periods by computing the probability of symptom onset and the distribution of incubation-period estimates across 1000 replicates for each dose.

## 3 Results

After deriving six two-scale within-host models, we simulated both the within-macrophage and within-host models. We began by obtaining new estimates for the rupture-size distribution across all six models. We ran the simulation for doses in the range of 1, 2, …, 500 *Legionella*. For each dose, we completed 1000 stochastic simulation iterations. Testing three rupture-size and two rupture-time models evaluated how macrophage variability affected dose-response predictions. This comparison not only validated our stochastic framework against [7], but also provided a methodology that could be applied to other intracellular pathogens to assess how cellular-level variability shapes population-level infection outcomes.

From these simulations, we recorded the proportion of iterations resulting in symptom onset as the probability of infection. For symptomatic individuals, the times at which symptom onset occurred were also reported. We then analysed the DR curves and estimated incubation-period distributions for each model. Finally, we conducted a sensitivity analysis by sampling parameters from the parameter estimate distributions to assess their impact on DR and incubation-period outcomes. In this sensitivity analysis, we analysed the Spearman correlation between parameter estimates and the outputted DR and incubation-period results.

### 3.1 Within-macrophage model

To explore how stochasticity within macrophages influences rupture-size distributions, we analysed the stochastic logistic growth birth–death (SLBD) process. In this section, we first present ensemble trajectories of the SLBD model. We then introduce three alternative implementations of rupture-size distributions, derived in equations (6), (7), and (8), and compare these with those proposed in [7]. Finally, we examine the impact of these approaches on the DR and incubation-period outputs, as well as the probability that a single surviving *Legionella* triggers symptom onset. To begin, we simulated the within-macrophage SLBD model and solved the master equations to complement these results (Figure 2).

**Fig 2.**
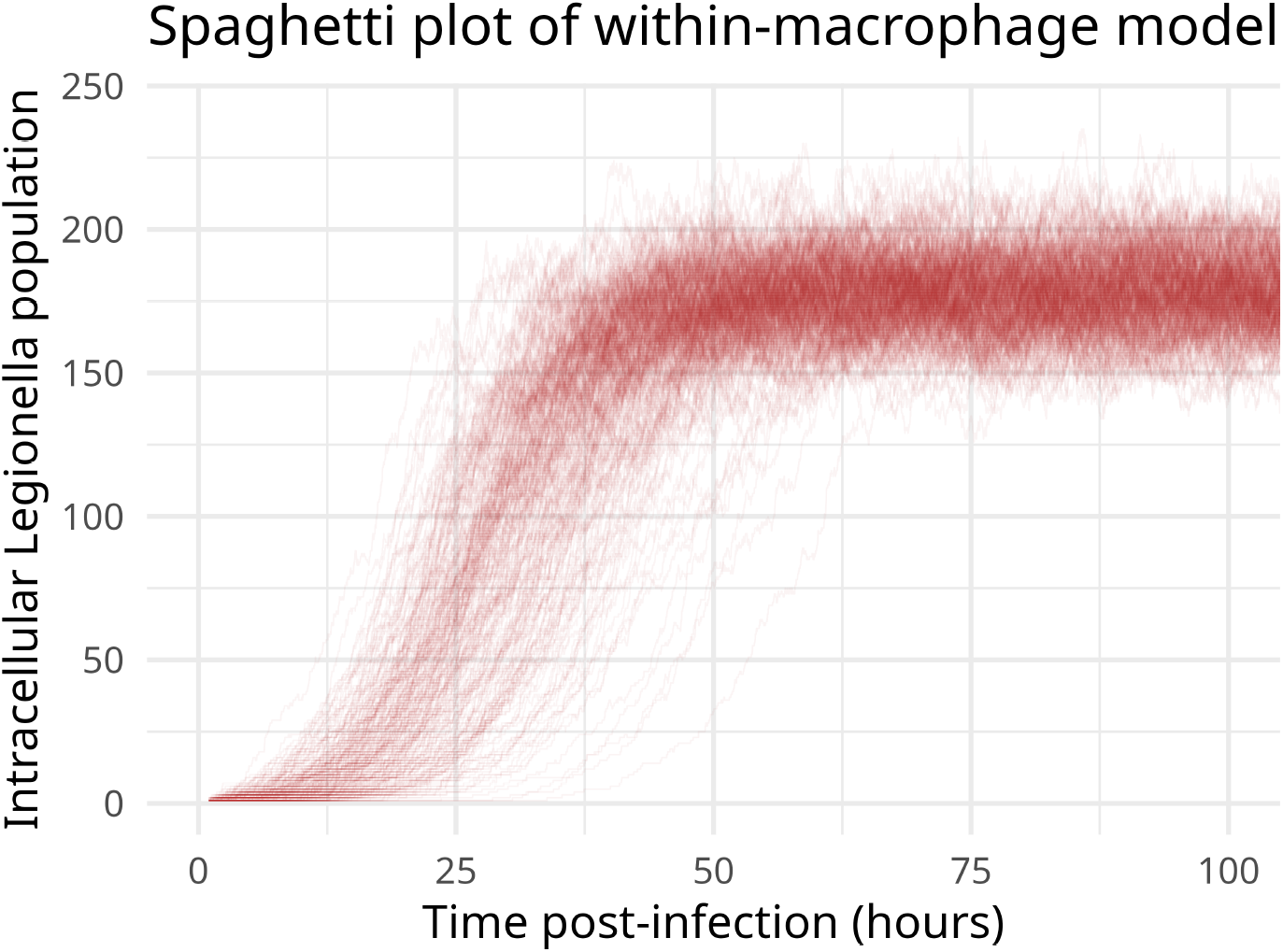
A plot of the within-macrophage SLBD model. The plot showed an ensemble trajectory plot of different realizations obtained from simulating the SLBD model master equations [equation (5)].

The general trajectories in Figure 2 approximated the deterministic logistic growth process given in equation (4). Our SLBD model was therefore a stochastic analogue of the earlier deterministic model, but it introduced variability at the macrophage level. In contrast, model C, which accounted for macrophage heterogeneity, assumed that within-macrophage dynamics remained deterministic. Additionally, models A and B assumed identical macrophages within the lungs. As illustrated in Figure 2, stochastic variability in bacterial replication rates among macrophages altered both rupture-size distributions and rupture timing. The master equation solution further highlighted how this variability translates into a probability mass distribution of outcomes.

Building on these results, we next combine the within-macrophage SLBD model with rupture-time distributions to generate rupture-size distributions. The resulting probability mass functions were given in equations (6), (7), and (8), and the distributions are visualised in Figure 3.

**Fig 3.**
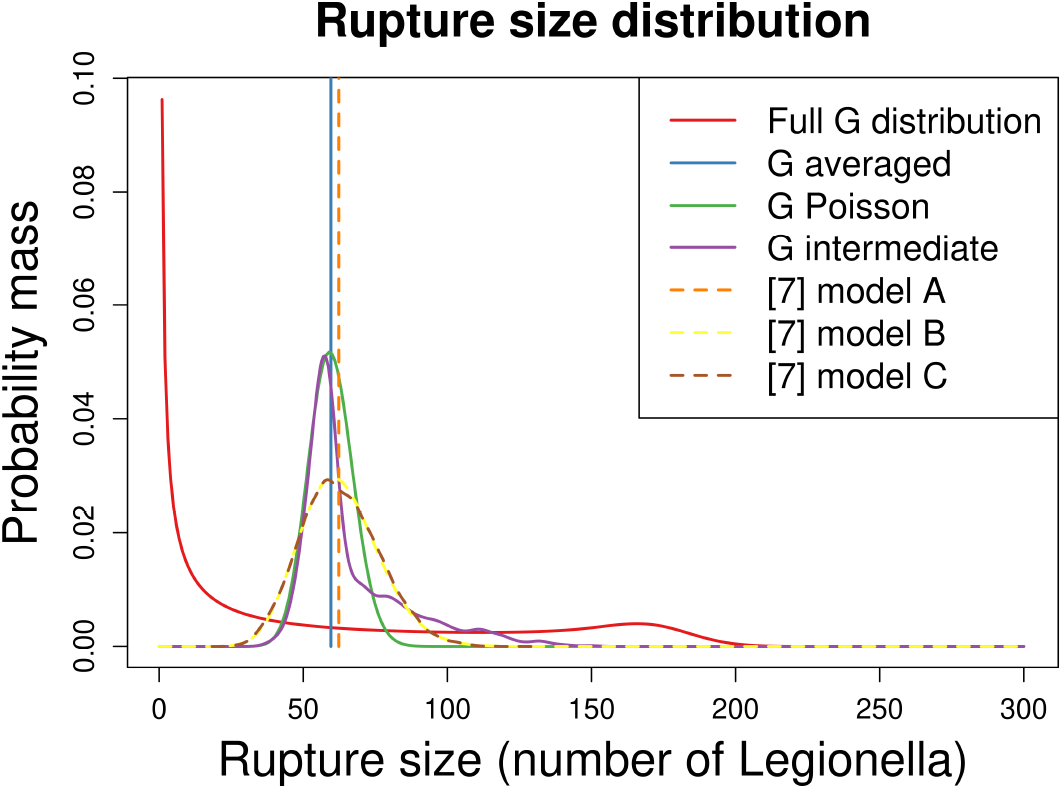
A plot of all six rupture-size distributions. Illustrated were the three rupture-size distributions derived in this study [average model 2_1_(*l*), intermediate model *q*_2_(*l*), and Poisson model *q*_3_(*l*)]. Further, the three rupture-size distributions from [7] were provided [point-estimate model A, negative binomial model B, and bootstrapped model C]. Additionally, the initial distribution of *G* across all rupture timings was provided.

As shown in red in Figure 3, the distribution of *Legionella* throughout an infected macrophage’s lifespan was skewed toward smaller sizes. The red curve is a bimodal distribution, which is caused by the fact that two possibilities occur for the stochastic within-macrophage model: some trajectories experience rapid early growth, while others remain near low population levels for some time before gradually increasing. Trajectories exhibiting rapid early growth reach the carrying capacity quickly, corresponding to the peak around *ψ Legionella*. In contrast, trajectories with slower initial growth may remain at low population levels for a time, creating a secondary mode at low bacterial counts. The combination of these two types of trajectories produces the observed bimodal distribution. This type of bimodal curve is also identified for *Francisella tularensis* [21] To address this, we considered three approaches to generating rupture-size distributions, corresponding to (6), (7), and (8). However, comparing the fully averaged model in equation (7) to the point-mass estimate in [7] yielded a larger point-mass estimate. Although the exponential rupture-time distribution underestimated the median rupture time compared with Erlang and Burr distributions, our stochastic SLBD model allowed for a possible slower bacterial growth in some realisations. As a result, macrophages that ruptured earlier released fewer bacteria than predicted by the deterministic logistic-growth model, reducing the median rupture size by approximately 2.74 *Legionella* compared with the exponential scenario. This difference produced a slightly lower mean rupture size overall.

Further, the Poisson and intermediate models provided modal values similar to those of models B and C. However, the intermediate model was more heavily weighted around its mean than models B and C, reflecting the lower variance of the Poisson distribution compared with the negative binomial (model B). At the same time, although models B and C had broader interquartile ranges and more dispersed densities around the mode, the intermediate model developed here spanned a larger overall range and was therefore more capable of generating large *G* values.

Finally, to connect these rupture-size distributions to potential simplifying interpretations for the DR relationship, we estimated the probability that a single surviving *Legionella* cell could trigger symptom onset. This probability was defined as:

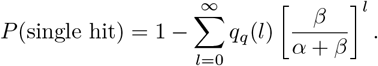

Here, the fraction *β/*(*α* + *β*) represents the probability that a single extracellular *Legionella* is eliminated by phagocytosis. Therefore, the summation gives the probability that *l* bacteria are released upon rupture of an infected macrophage and that all *l* progeny are subsequently removed. Using this expression, we obtained probabilities of a single hit hypothesis (i.e., a single *Legionella* surviving phagocytosis is sufficient for eventual symptom onset) estimated as 99.08%, 99.07% and 98.89% for the three rupture-size models (*q*_1_(*l*), *q*_2_(*l*), and *q*_3_(*l*), respectively). Equivalently, the probability that a single surviving *Legionella* did not result in eventual symptom onset was 0.92%, 0.93% and 1.11% for the three models, respectively. Although a single survival of *Legionella* during phagocytosis and eventual macrophage rupture is not sufficient to guarantee eventual symptom onset, these results indicated that a single-hit hypothesis provided a reasonable description of the DR relationship for Legionnaires’ disease.

### 3.2 Dose–response results from the two-scale within-host simulations

To assess the impact of rupture assumptions on infection risk, we fitted DR models to the simulated data from all six combinations of rupture-size and rupture-time distributions. Because our earlier analysis estimated a ≥ 98.89% probability that a single surviving *Legionella* leads to symptom onset, we applied the exponential DR model, which was based on the single-hit hypothesis. Fitting this model allowed us to estimate the infectious dose required to cause illness in 50% of individuals (ID_50_). We summarize the estimates of ID_50_ for the six two-scale models considered here, as well as for the three within-host models A, B and C (Figure 4).

**Fig 4.**
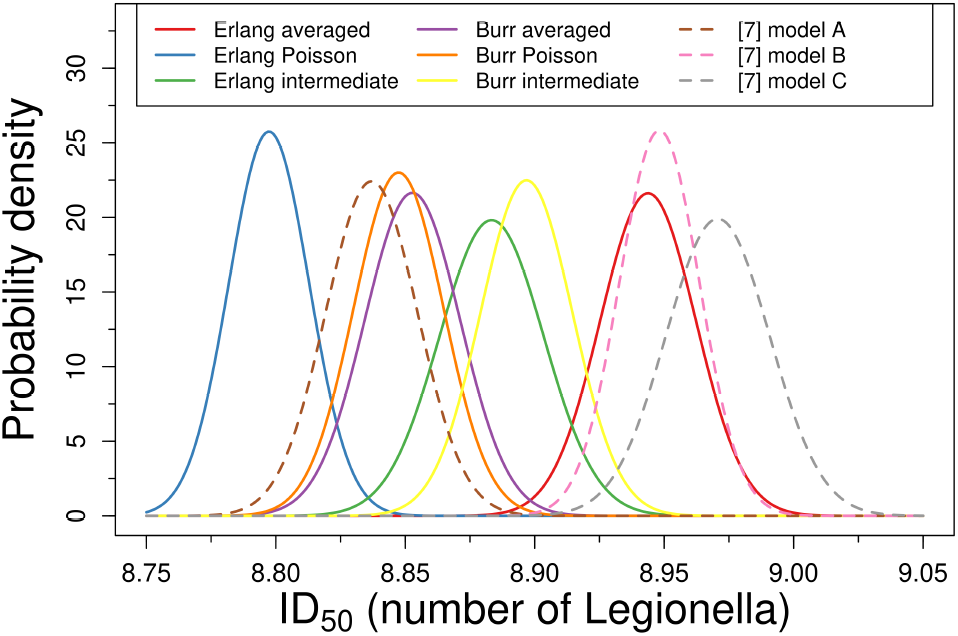
Estimates of the ID_50_ from fitting the exponential DR model to the dose–response results from each of the six two-scale models. For comparison, estimates of the ID_50_ from fitting the exponential DR model to previous three within-host models A, B and C.

The ID_50_ estimates range narrowly between 8.79 and 8.94, indicating that differences among models, though statistically significant, remain small. This observation suggests that although assumptions about rupture timing and averaging can subtly influence infection probability, the overall public health risk may be largely robust to these modeling choices. In addition, the ID_50_ estimates (8.75–9.05) do not display a consistent pattern with respect to rupture-time or rupture-size assumptions, and the small discrepancies are comparable to the stochastic variation inherent in the simulations. These findings raise the possibility that robustness to rupture assumptions may be a general feature of the DR relationship of stochastic infection dynamics, a hypothesis that could be further tested by comparison with other intracellular pathogens.

To illustrate these results, we provide plots of the infection probability against deposited dose. The six simulated DR curves overlap closely, reinforcing the observation that ID_50_ estimates differed only marginally across models (Figure 5).

**Fig 5.**
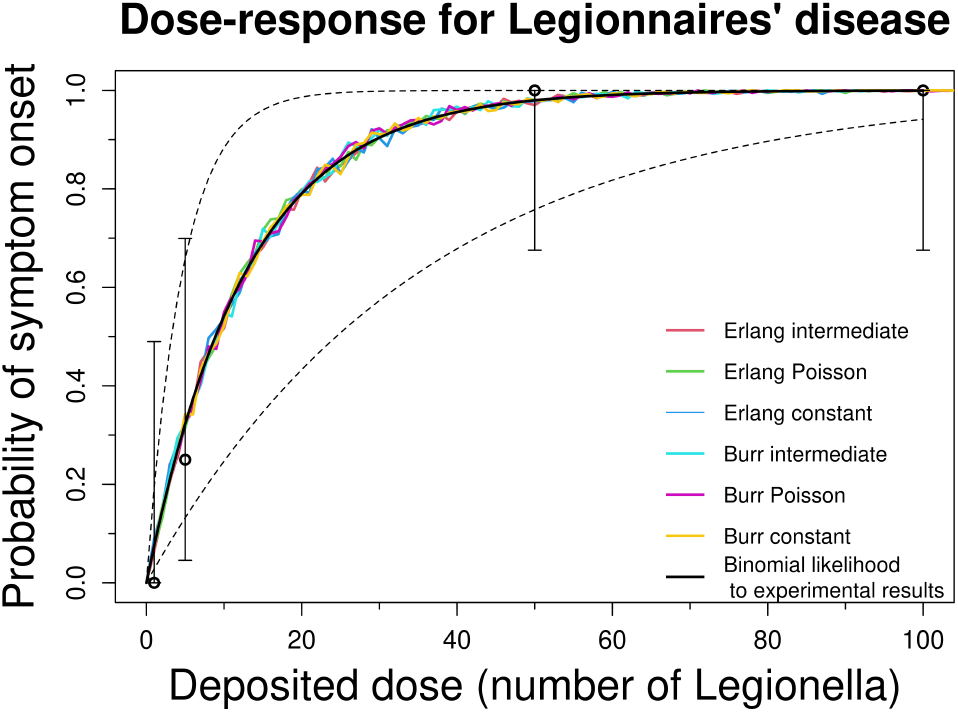
A plot of the guinea pig DR data as four data points representing the proportion of guinea pigs that developed symptoms [20]. Error bars were provided for each point from the [20] dataset to provide the uncertainty in their values when used to model the probability of onset of symptoms in this experiment. An exponential DR model was fitted to the data in [20] using a binomial likelihood approach. The DR data obtained from simulating the three within-host models were presented for comparison.

Finally, our simulated DR curves closely reproduced the experimental data. The exponential DR model fitted directly to the experimental data yielded an ID_50_ estimate consistent with our simulations, ranging from 8.79–8.94 *Legionella*. This agreement demonstrated that the two-scale models developed here can successfully recapitulate the experimentally observed DR relationship of Legionnaires’ disease, in line with results reported previously.

In [7], we estimated an ID50 of (3.22, 24.44) from the DR experiment in [20]. This confidence interval is wide, reflecting the small sample size in [20], where the probability of symptom onset was estimated from a proportion of eight individuals. In contrast, the ID50 confidence intervals estimated in this paper are much narrower, reflecting the larger sample size (1000) used at each dose level (1–500). A wider confidence interval would likely be obtained if population heterogeneity, as considered in [7], were incorporated. However, we chose not to include this here, as our focus was specifically on the effects of updating the rupture-time distribution. Notably, the confidence intervals are non-overlapping for some of the six models developed in this chapter, highlighting the sensitivity of the ID50 to model assumptions. Nevertheless, because the confidence intervals are narrow and close in magnitude (despite being non-overlapping), the resulting DR curves remain relatively robust to these variations in ID50.

### 3.3 Incubation-period results from the two-scale within-host simulations

We next analyse dose-dependent incubation periods estimated using our six within-host models. For each simulation, we recorded the incubation period of symptomatic individuals across deposited doses, generating a dose-dependent incubation-period dataset. From this dataset, we estimate the mean, median, standard deviation, skewness, and kurtosis, and compare these results to the models A, B and C. We summarize these comparisons below (Figure 6).

**Fig 6.**
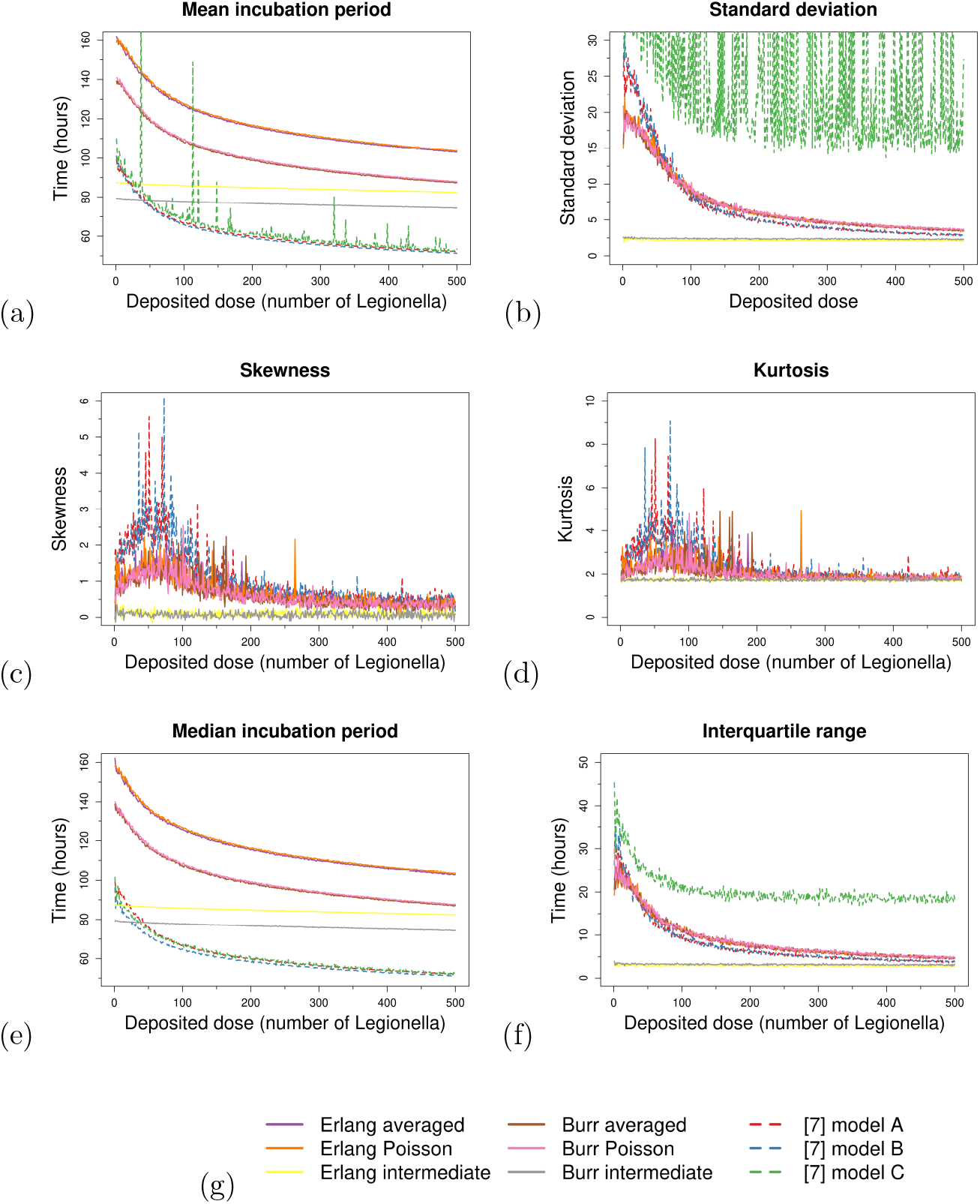
Moments of the dose-dependent incubation period for each of the six two-scale within-host models and the three models A, B, and C. Sub-figures (a–b) show the mean and standard deviation, respectively. Sub-figures (c–d) show skewness and kurtosis, respectively. Sub-figure (e) shows the median across doses. Sub-figure (f) shows the interquartile ranges across doses. Sub-figure (g) presents the legend that is shared across all sub-figures.

For both Erlang and Burr rupture-time distributions, the intermediate model yields a consistent mean and median across the full dose range (1–500). These distributions also produce stable standard deviations and approximately normally distributed behaviour (skewness near 0, kurtosis near 3). Rupture variability therefore dominates incubation-period timing, outweighing the effect of initial dose on eventual deterministic growth. The Burr distribution produces consistently shorter incubation-period estimates than the Erlang distribution, likely because its slightly larger median rupture time releases more *Legionella* per rupture, accelerating the approach to the threshold *T*_*L*_. Across doses, the Burr results remain roughly parallel to the Erlang results but are shifted earlier by about 20 hours.

Moreover, for each rupture-time distribution, the average model yields slightly lower incubation-period estimates than the Poisson model. This observation is an artefact of the fact that the Poisson model occasionally generates large rupture events that accelerated the process towards deterministic growth [8] relative to the averaged model. In contrast, the intermediate model shows less dependence on deposited dose, with its distribution more heavily influenced by rupture-size variability. This explains why correlations between dose and incubation period are weaker under the intermediate assumption (see discussion on Spearman correlation below).

Additionally, Figure 6(c–d) provides validation to work in [12], which had shown that the gamma, Weibull and lognormal distributions are insufficient as an incubation-period model for Legionnaires’ disease. Only by considering a distribution for the (non-dose-dependent) incubation period that provides more flexibility in the skewness and kurtosis, such as the Burr distribution, could one obtain reliable incubation-period modelling results [12].

Furthermore, models A, B and C predict much shorter incubation periods. By incorporating stochastic within-macrophage growth and rupture timing, our six models produce median incubation periods of 103.5–159.0 hours across doses 1–500. These results are consistent with experimental observations and correct the 50.5–73.1 hour underestimation in [7]. This highlights that deterministic models systematically underestimate incubation periods by neglecting stochastic macrophage-level processes.

Beyond this, we note that model C in Figure 6(a) exhibits spikes in the mean incubation period, whereas the median incubation period for model C in Figure 6(e) remains smooth. This occurs because model C samples new parameter estimates in each simulation; with the number of simulations reported in [7], the mean has not fully converged to the true value, and extreme incubation-period estimates produce the observed spikes. In contrast, the median is robust to such extreme values. Furthermore, in Figure 6(f), model C displays a larger interquartile range than the new models developed in this paper. While our new within-host models correct the mean and median estimates, removing bias and consistently aligning with observed human outbreak data, realistic variance is best captured by sampling from the full parameter distributions rather than using only the central estimate in each simulation. Therefore, combining the methodology introduced in this paper with model C’s approach of sampling from parameter distributions provides a framework that accurately captures both the mean and variance of the incubation period for LD.

Finally, to quantify dose–incubation relationships, we calculate Spearman correlations between dose and incubation period. Results range from *τ* = – 0.467 (Burr–intermediate) to *τ* = – 0.865 (Erlang–averaged and Erlang–Poisson). Across all six models, we find strong negative correlations, indicating that higher deposited doses shortened incubation periods. The Burr models consistently yield weaker correlations than the Erlang models, and the weakest correlations occurs under the intermediate rupture-size distribution. This shows that rupture-size variability can mask the expected dose–incubation relationship, implying that heterogeneous environments may reduce the predictive value of dose-based risk assessments.

Overall, stochastic rupture timing shifts incubation-period estimates by 49–64% compared with deterministic assumptions, underscoring its critical role in host response. Errors in rupture-time models propagate across successive events, biasing incubation-period estimates by hours or even days. Moreover, allowing rupture-size distributions to depend on time amplifies this effect: incorrect timing assumptions underestimate rupture-size distributions, altering the trajectory to deterministic exponential growth. For example, an exponential rupture-time distribution underestimates incubation periods by several hours, leading to biases of 49–64%. These results suggest that within-host models assuming Markovian macrophage dynamics may systematically underestimate early case onset.

### 3.4 Sensitivity analysis

We conduct a sensitivity analysis to assess how variation in model parameters influences DR and incubation-period estimates. The analysis is performed at a deposited dose of nine *Legionella*, which was the closest integer to the estimated ID_50_ across all models. For each parameter, we sample values from its distribution while holding non-dependent parameters fixed. Dependent parameters are recalculated as point estimates to avoid bias. For each parameter sample, 1000 within-host simulations are performed to record the probability of symptom onset (as a proxy for infection risk) and the incubation-period estimates for symptomatic cases. This procedure is repeated for 100 samples of each parameter, and Spearman correlations are calculated to quantify monotonic relationships between parameter values and outcomes (Figure 7).

**Fig 7.**
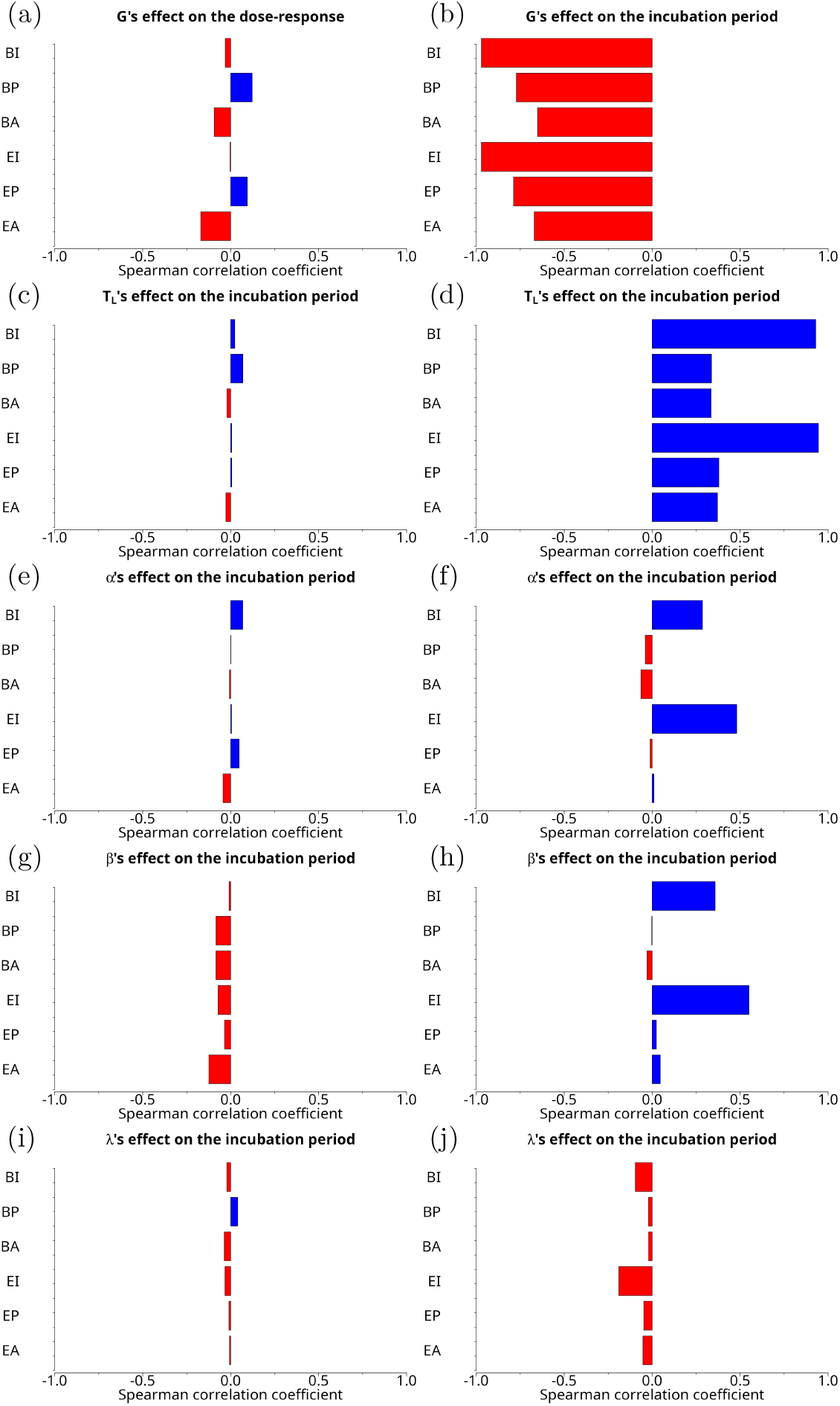
Plots of the sensitivity analysis for the six two-scale within-host models. In this plot, the abbreviations are as follows: *BI* (Burr–intermediate), *BP* (Burr–Poisson), *BA* (Burr–average), *EI* (Erlang–intermediate), *EP* (Erlang–Poisson), and *EA* (Erlang–average). Subfigures (a) and (b) show the DR and incubation period results, respectively, from the *G*-varying simulation. Subfigures (c) and (d) show the DR and incubation period results, respectively, from the *T*_*L*_-varying simulation. Subfigures (e) and (f) show the DR and incubation period results, respectively, from the *α*-varying simulation. Subfigures (g) and (h) show the DR and incubation period results, respectively, from the *β*-varying simulation. Subfigures (i) and (j) show the DR and incubation period results, respectively, from the *λ*-varying simulation.

Across all six models, varying parameters have only weak effects on the DR relationship. This indicates that infection probability is relatively robust to parameter uncertainty. In contrast, incubation periods show marked sensitivity to several parameters. For all models, increasing the mean rupture size consistently shorten incubation periods, as larger ruptures accelerate extracellular *Legionella* accumulation and reduce time to threshold *T*_*L*_. The intermediate models illustrate this effect most strongly, with mean incubation periods ranging from 98–223 hours depending on rupture size. Further, phagocytosis parameters (*α* and *β*) and median rupture time (*λ*) have little effect under either the Erlang or Burr models with averaged or Poisson rupture-size assumptions. However, under the intermediate rupture-size model, increasing *α* or *β* lengthened incubation periods. Faster phagocytosis relative to rupture events deplete extracellular populations more rapidly, delaying symptom onset. Similarly, decreasing *λ* (slowing rupture events) increases incubation times by enhancing phagocytic clearance. In these cases, extracellular populations could vary by up to 179-times between successive rupture events. Threshold *T*_*L*_ of *Legionella* also have a strong effect, especially in intermediate models, where Spearman correlations exceeded 0.8. Higher thresholds substantially prolong incubation periods, consistent with the role of *T*_*L*_ in defining the symptomatic transition.

Overall, the sensitivity analysis highlights that incubation-period predictions are most affected by mean rupture size, phagocytosis rates, and rupture timing. In contrast, probability of symptom onset is comparatively insensitive to parameter variation. These findings suggest that variation in modelled host immune parameters, particularly phagocytosis rates and macrophage rupture dynamics, could significantly affect the timing of disease onset. Importantly, our approach produces a lower bound on incubation-period estimates that was consistent with experimental observations ( ≈ two days). This reinforces the fact that stochastic macrophage-level processes must be incorporated to accurately model infection dynamics. Future experiments that manipulate rupture size or phagocytosis rates could directly test these predictions and extend them to other intracellular pathogens.

## 4 Discussion

This study developed models to extend the work in [7] to address specific limitations in the within-host model of Legionnaires’ disease, particularly regarding rupture-time distributions and within-macrophage variability. Three principal extensions were introduced: (1) replacing the deterministic within-macrophage model with a stochastic analogue, (2) introducing non-Markovian distributions for macrophage rupture timing, and (3) developing formal methods for modelling time-dependent rupture-size distributions. The stochastic model allowed within-macrophage dynamics to vary even among infected macrophages sharing identical parameters, whereas the previous model varied only macrophage capabilities. For rupture timing, Erlang and Burr distributions were implemented to better fit observed data, and a non-Markovian Gillespie algorithm was employed for simulation. By combining these elements, we simulated DR curves and dose-dependent incubation periods, and compared results with the exponential DR model and previous rupture-time distributions reported.

The previous model (discussed in Section 2(b)) underestimated the mean and variance of incubation periods when assuming either within-host randomness or population heterogeneity alone (models A and B); only by assuming both did variance partially match human data (model C). To address this limitation, our model focused on within-host stochasticity and incorporated modified rupture-timing and rupture-size distributions. This approach enabled us to assess whether the model’s limitations in representing rupture events were responsible for the underestimation of both the mean and the variance. Consequently, simulations of the average and Poisson rupture-size models showed that, for exposure doses typical of Legionnaires’ disease outbreaks, the mean onset of illness occurred five–six days post exposure. This outcome quantitatively aligned with outbreak data [12, 28, 29] and corrected the 50.5–73.1 hour discrepancy reported in [7]. Simulated incubation-period distributions ranged from 2–13 days, primarily 2–10 days, which was consistent with observed human outbreaks [12].

Our model provides more accurate predictions of both incubation-period distributions and DR curves than the previous Markovian framework. Specifically, the estimated ID_50_ ranged between 8.79–8.94 *Legionella* across models, which is consistent with experimental data [7, 20]. This small range indicates that the DR results are fairly robust to small ID_50_ variation resulting from different rupture-size models. Although DR analysis remained consistent with the prior framework, our modifications extend the model to reproduce published human incubation-period data and address earlier limitations arising from the Markov rupture-time model. Additionally, biological processes such as cytokine-mediated inflammation or recruitment of additional immune cells were not incorporated, and phagocytic cell recruitment to the lungs and the adaptive immune response were excluded. Nonetheless, the innate immune response model accurately reproduced DR and incubation-period dynamics, indicating that the adaptive immune response primarily affects pathogen clearance rather than symptom onset probability and timing.

Regarding the within-host rupture-time models, the events within models A, B and C were all modelled using exponential distributions. Although reasonable for phagocytosis due to random mass-action bacterial movement assumptions, exponential distributions poorly fit macrophage-rupture timing. This choice, necessary for a continuous-time Markov chain, constrained the model’s ability to reproduce observed incubation-period distributions. By employing Erlang and Burr distributions, the median rupture time increased relative to the exponential distribution, producing longer incubation periods after accumulated rupture events. Consequently, our model predicted incubation periods within the observed human range, and the adoption of non-Markovian distributions resolved this limitation whereas DR predictions remained consistent with the Markovian models.

This work provides an improvement over the CTMC-based approaches presented in [9, 10]. While developing a CTMC using an Erlang distribution with the method of stages offers a parsimonious and analytically tractable solution for describing rupture-time distributions, it may be overly simplistic. In particular, the Erlang distribution does not adequately capture the tails of the empirical rupture-time data [11], a limitation addressed by the Burr nMGA model. An alternative approach could employ a phase-type distribution, such as the hypoexponential, which would allow for a Markovian formulation potentially as accurate as the Burr model. This approach has been considered for *Francisella tularensis* and *Bacillus anthracis* [21–23]. However, the Burr model provides a high level of flexibility, fitting the observed rupture-time data extremely well, and it integrates naturally into the nMGA algorithm. Any potential gains in accuracy from using a phase-type model would likely be outweighed by the increased computational cost associated with its implementation, making the Burr nMGA approach a practical and efficient choice for capturing the rupture-time dynamics.

Furthermore, the results from simulating the model in Section 2(b) suggest that animal data might be insufficient for developing a within-host model [7]. By identifying the Markov assumption on rupture timing as the primary limitation, we implemented stochastic macrophage dynamics and non-Markovian rupture timing, demonstrating that animal data did not compromise result reliability. Specifically, the hypothesis that lacking an adaptive immune response model caused lower incubation periods appeared inaccurate. Including the adaptive immune response may later add further mechanisms that kill *Legionella*. However, at least five days are required for effects from the adaptive immune response [24, 25]. By this stage, symptom onset has likely occurred, or bacterial growth has reached a phase of near-deterministic exponential expansion, at which point the adaptive immune response has minimal affect on reducing bacterial replication rates. These hypotheses indicate that the adaptive immune response does not critically influence symptom onset timing or probability, but remains essential for modelling pathogen clearance later in infection.

In conclusion, this study presented the second mathematical within-host model of Legionnaires’ disease and was the first to incorporate non-Markovian dynamics while addressing prior limitations. As a result, DR curve and incubation-period predictions quantitatively aligned with observed experimental and outbreak data. Moreover, mechanistic modelling of within-macrophage and rupture-timing dynamics allowed a rigorous evaluation of the single-hit hypothesis, which we showed does not fully apply to *Legionella* infection, although it provided a reasonable approximation (with a probability ≥ 98.89% chance of a single hit resulting in symptom onset). Beyond Legionnaires’ disease, this framework can be applied to other intracellular pathogens, including *Francisella tularensis* and *Coxiella burnetii*, as these pathogens exhibit similar within-host and within-macrophage dynamics. The model can also be extended to incorporate additional immune cells, such as dendritic cells, monocytes, and neutrophils, during later stages of infection. Finally, this modelling approach has direct practical utility for public health: improved incubation-period predictions support source identification in outbreak investigations, whereas enhanced DR understanding provides quantitative guidance for environmental risk assessment.

